# Antibiotic resistance evasion is explained by rare mutation frequency and not by lack of compensatory mechanisms

**DOI:** 10.1101/374215

**Authors:** Dimitrios Evangelopoulos, Gareth A. Prosser, Angela Rodgers, Belinda M. Dagg, Bhagwati Khatri, Mei Mei Ho, Maximiliano G. Gutierrez, Teresa Cortes, Luiz Pedro S. de Carvalho

## Abstract

Drug resistant infections represent one of the most challenging medical problems of our time. D-cycloserine is an antibiotic used for decades without appearance and dissemination of antibiotic resistant strains, making it an ideal model compound to understand what drives resistance evasion. We investigated why *Mycobacterium tuberculosis* fails to become resistant to D-cycloserine. To address this question we employed a combination of bacterial genetics, genomics, biochemistry and fitness analysis *in vitro*, in macrophages and in mice. Altogether, our results suggest that the ultra-low mutation frequency associated with D-cycloserine resistance is the dominant factor delaying the appearance of clinical resistance to this antibiotic in bacteria infecting humans, and not lack of potential compensatory mechanisms.

**One Sentence Summary:** We show that the lack of D-cycloserine resistance in *Mycobacterium tuberculosis* is due its ultra-low mutation frequency rather than lack of compensatory mechanisms.

---

The number of human infections caused by bacteria resistant to antibiotics has dramatically increased in the last decades (*1-3*). Currently, almost all drug classes in clinical use are matched by evolved mechanisms of resistance in bacteria (*4, 5*). Some of these bacteria are pan-resistant and cannot be treated with any approved drugs (*6*). While we understand very well how these antibiotics work and how antibiotic resistance works in many cases, we still fail to recognize how some drugs select for resistant in months while others need several decades (resistance evasion)(*7, 8*).

As for all other human pathogens, *Mycobacterium tuberculosis*, the etiological agent of human tuberculosis (TB), has acquired mutations in its genome and evolved resistance mechanisms to almost all drugs that it has encountered during chemotherapy (*9*). Currently, the genetic basis for first- and second-line antibiotic resistance in *M. tuberculosis* is well characterized, with one exception, the antibiotic D-cycloserine (DCS) (*10*). DCS belongs to the core second line treatment group C listed by WHO guidelines for treatment of multi-drug and extensively-drug resistant-TB (MDR/XDR-TB). Given its activity and lack of reported resistance in strains infecting humans, DCS has been called “the cornerstone option” for treating drug resistant TB cases (*11*). This feature makes DCS the only antibiotic that has been used in humans for almost seven decades that has evaded resistance selection in bacterial populations (*12, 13*). From a practical standpoint, the molecular and cellular determinants for DCS resistance-evasion are highly attractive features that if understood could be incorporated into antibiotic drug discovery programs aimed at developing superior antimicrobial agents for the treatment of tuberculosis and other infectious diseases.

In contrast to the mechanisms of resistance, the mechanism of DCS action has been well studied since its discovery in the 1950’s. In all bacteria tested, DCS inhibits two enzymes of the D-Ala-D-Ala branch of peptidoglycan biosynthesis: alanine racemase (Alr) and D-Ala:D-Ala ligase (Ddl) (*14, 15*). In *M. tuberculosis*, it has been established that despite DCS still inhibiting both targets, the principal mechanism of bacterial death is inhibition of D-Ala:D-Ala ligase (encoded by the *ddlA* gene), instead of inhibition of alanine racemase (encoded by the *alr* gene) (*16, 17*). This conclusion is supported by earlier independent work which demonstrates a high catalytic excess of alanine racemase in *Mycobacterium smegmatis* (*18*). Surprisingly, until very recently there were no well-defined, associated mutations conferring DCS resistance in the clinic. It has been described that in the fast-grower, free-living *Mycobacterium smegmatis*, artificial overexpression of both Alr and DdlA result in DCS resistance (*19, 20*). Furthermore, a point mutation in the *cycA* gene, encoding for the DCS transporter, is responsible for the natural DCS resistance of *Mycobacterium bovis* BCG (*21*). Recently, whole-genome sequencing (WGS) data from clinical isolates including MDR and XDR-TB strains identified non-synonymous single nucleotide polymorphisms (SNPs) in the *alr* gene with the possibility of some of them conferring low-level DCS resistance (*22-24*). However, whether these SNPs are relevant during TB treatment remains unclear.

To identify possible indications of DCS resistance in *M. tuberculosis*, we first analysed the distribution and frequency of SNPs in the DCS target genes *alr, ddlA, cycA* as well as the *ald* gene, as it has been recently implicated in low-level DCS resistance (*24*). We screened a published dataset comprising 1,601 genomes of *M. tuberculosis* clinical isolates (*25*) and compared it to the polymorphisms found in *rpoB* and *gyrA*, associated with drug resistance to rifampicin (*26*) and fluoroquinolones (*27*), respectively. We identified scarce non-synonymous SNPs in the *ddlA* gene (0.80 %) and on *alr* gene (0.98 %) and a slightly higher frequency of non-synonymous SNPs in the, *cycA* (1.26 %) and *ald* genes (1.52 %) when compared to the *gyrA* and *rpoB* (1.99 and 1.85 %, respectively) (Fig. S1 and table S1). Therefore, these results demonstrate a low degree of SNPs in the genes encoding DCS targets, in agreement with lack of DCS resistance in clinical isolates.

To investigate the biological causes of such low resistance frequency in *M. tuberculosis*, we examined the frequency of spontaneous mutations conferring resistance to DCS, by carrying out fluctuation analysis experiments (*28*). Fluctuation analysis indicates that the frequency of spontaneous mutations conferring DCS resistance in *M. tuberculosis* is ultra-low (10^-11^). This result is in agreement with the only other report of the mutation rate of DCS with *M. tuberculosis* (*29*). This ultra-low frequency is consistent with DCS engaging two lethal targets in *M. tuberculosis*, Alr and Ddl, and represents an important barrier for the selection of resistance. Nevertheless, under antibiotic pressure this frequency might be different, as has been suggested from a recent study reporting DCS clinical resistance (*22, 24*).

To investigate the mechanisms responsible for DCS resistance in *M. tuberculosis*, we isolated and characterized spontaneous DCS-resistant mutants (DCS^R^), obtained independently. Eleven DCS^R^ strains were isolated and whole genome sequenced along with the parental *M. tuberculosis* H37Rv strain (table S2). A total of 85 non-redundant SNPs were identified across the resistant isolates, when compared against the reference genome of *M. tuberculosis* H37Rv. After excluding SNPs within repetitive regions, a total of eleven SNPs were targeted for further analysis (Fig. 1A and table S3). Among the genes known to be potential DCS targets or associated with DCS resistance, we only detected mutations in the *alr* gene (Fig. 1A and B). Eight out of the eleven mutants shared a non-synonymous SNPs in position 3840391, creating an aspartate (D) to asparagine (N) substitution at codon 322 of the *alr* gene. Furthermore, DCS^R^8 and DCS^R^11 shared a mutation located within the promoter region (C 3841405 A) of the *alr* gene. This mutation changes the −10 promoter region from a non-TANNNT −10 motif to an extended −10 promoter consensus sequence (*30*), suggesting possible transcriptional changes. In addition, we detected a 142 bp deletion upstream of the *alr* promoter in DCS^R^5, possibly in a region where a transcriptional regulator might bind. *In silico* analysis of transcriptional factor binding site indicates that no known transcriptional regulators bind to this region, suggesting it could be a novel regulator. After screening of these two SNPs in the Coll *et al*., (*25*) dataset, only a single occurrence of the first SNP (C 3840391 T; D322N) was reported in a Russian strain belonging to lineage 1. Based on the pattern of non-synonymous SNPs detected among the mutant strains, we grouped them into six different groups (Fig. 1A and B). Furthermore, mutations in the promoter or repressor of *alr* seem to have evolved in parallel, as all the secondary mutations found in these strains are also present in the *alr* D322N mutant strains. These data highlight the central role of *alr* in DCS resistance in *M. tuberculosis*.

**Fig. 1.**
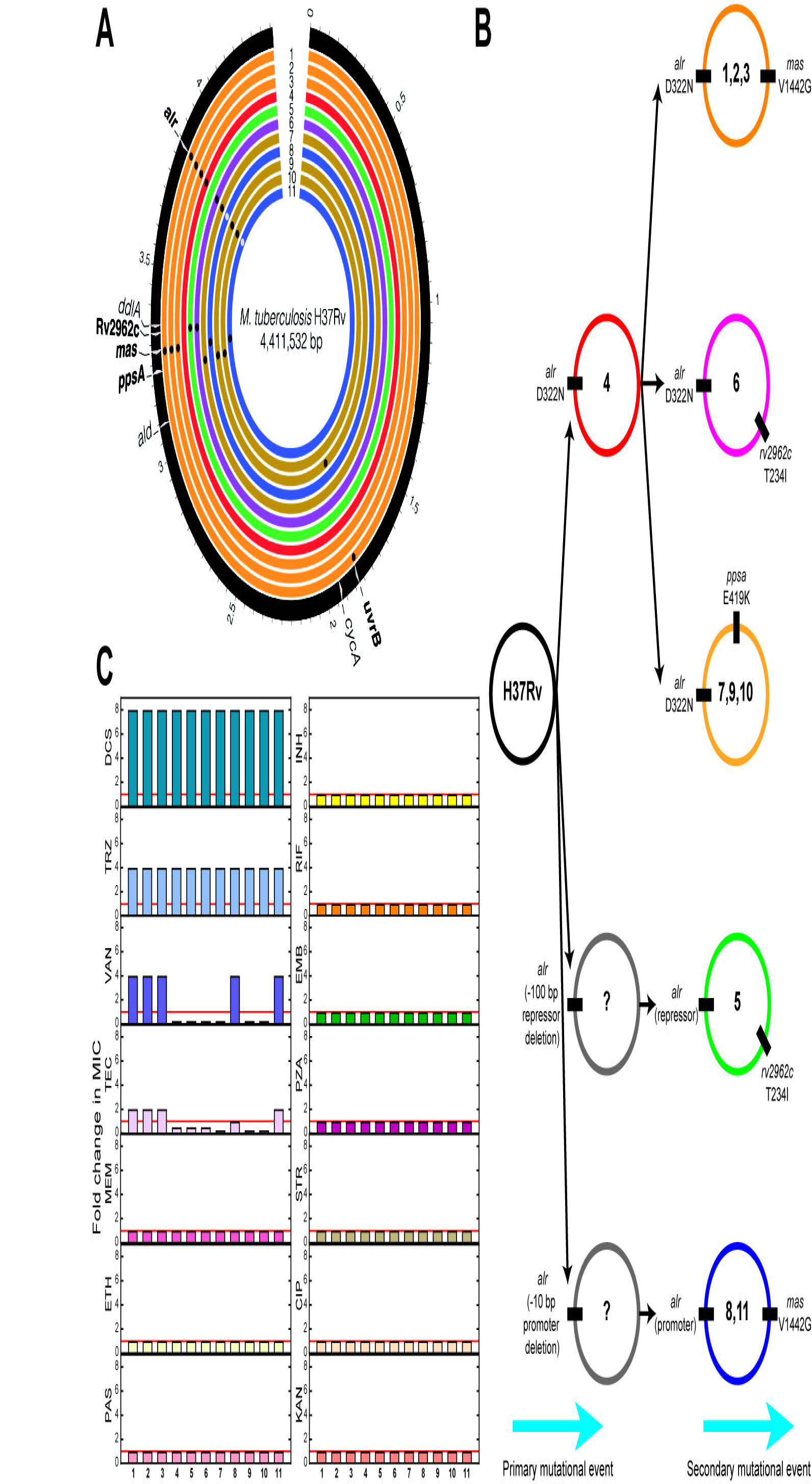
Distribution of DCS resistant mutations, evolution of resistant strains and drug susceptibility profile of the DCS^R^ strains. (**A**) Genome visualization illustrating the SNPs identified in the DCS^R^ strains compared to the parent, DCS^S^ strain. Moving from the outer to innermost ring, genome reference is indicated in black, with genes harboring SNPs or representing DCS targets highlighted in white, followed by DCS^R^ strains 1 to 11. Presence of non-synonymous SNPs is represented with black dots whilst SNPs within the promoter region of the *alr* gene are colored in grey. Names of genes with identified SNPs have been highlighted in bold. Mutants have been color-coded, based on the types and polymorphisms observed in the genome. Circular map was generated using Circos (Krzywinski et al., 2009). (**B**) Schematic representation of evolution of DCS resistance. Circles represent the genomic DNA of each strain and numbers refer to the specific mutant ID. (**C**) Drug susceptibility pattern of DCS resistant mutants (MIC_90_ fold change DCS^R^/parent strain). Drug abbreviations used: DCS; D-cycloserine, TRZ; terizidone, VAN; vancomycin, TEC; teicoplanin, MEM; meropenem, INH, isoniazid, RIF, rifampicin, EMB; ethambutol, PZA; pyrazinamide, STR; streptomycin, CIP; ciprofloxacin, KAN; kanamycin, ETH, ethionamide, PAS; para-aminosalicylic acid, MTZ; metronidazole. The MIC_90_ of the parent, DCS^S^ strain was 6.25 μg/mL. Data are representative of three independent experiments.

We next tested the extent of resistance and whether these mutants display any cross-resistance against other antitubercular drugs. Minimal inhibitory concentrations (MIC_90_) were evaluated for all 1^st^ line and 2^nd^ line antitubercular drugs, as well as other peptidoglycan targeting drugs. All eleven mutants are 8-fold more resistant to DCS when compared to the parent strain (Fig. 1C). In line with these results, all eleven mutants were also 4-fold more resistant to terizidone (TRZ), a better tolerated DCS pro-drug that contains two equivalents of DCS per molecule (*31*). Interestingly, these mutants showed differential susceptibility patterns when tested against vancomycin (VAN) and teicoplanin (TEC). VAN and TEC are glycopeptide antibiotics that bind to the terminal D-Ala:D-Ala moiety of the pentapeptide stem of peptidoglycan and prevent the crosslink of the peptidoglycan chains. Mutants DCS^R^1, DCS^R^2, DCS^R^3, DCS^R^8 and DCS^R^11 were four-fold more resistance to vancomycin whereas the rest of the DCS^R^ mutants were four-fold more susceptible when comparing to parent strain (Fig. 1C). It seems that the DCS^R^ strains containing the *mas* gene (V1442G) SNP are more resistant to VAN and TEC than the other strains. The *mas* gene encodes a polyketide synthase involved in mycocerosic acid synthesis, a part of the highly lipophilic outer membrane lipid phthiocerol dimycocerosate (pDIM). In line with our results, changes in pDIMs have been previously shown to affect the susceptibility of *M. tuberculosis* strains to VAN (*32*). Importantly, not all DCS^R^ mutants have altered pDIMs (Fig. S2). The sensitivity to glycopeptides, triggered by resistance to DCS also highlights the collateral implications that certain antibiotic resistant strains may have with other antibiotics (*33*). Of broad interest, the fact that a single SNP can confer VAN resistance in *M. tuberculosis* is highly unusual. For example, in glycopeptide-resistant enterococci, a series of mobile elements containing at least four but up to seven genes is required to cause VAN resistance (*34, 35*). Importantly, no cross-resistance to any 1^st^ line and 2^nd^ line drugs was observed in the 11 DCS^R^ mutants (Fig. 1A), indicating that these strains bear mutations that are mechanistically linked to DCS resistance, instead of general effects such as changing proton motive force or antibiotic efflux.

We next sought to investigate the potential fitness cost that these mutations might confer, which if high, could explain the scarcity of DCS resistant *M. tuberculosis* strains in the population. Given the slow-growing nature of *M. tuberculosis,* we carried out these experiments as a function of time, by measuring fitness kinetics. Kinetic analysis of fitness cost also precludes interpretation of single time points, which might not reflect the true phenotype. We examined fitness *in vitro,* in standard Middlebrook 7H9 medium, using a direct competition assay; inside interferon-gamma-stimulated or naïve human monocyte-derived macrophages; and finally, *in vivo* in a mouse model of infection (Fig. 2). *In vitro*, DCS^R^4, the mutant that solely contains the primary mutation D322N in Alr exhibited a transient fitness cost (days 7 and 14) which was lost by the end of the assay (day 56) (Fig. 2A). Similar fitness kinetics was also observed with DCS^R^5, the mutant having a deletion upstream of the *alr* gene. However, the occurrence of the *mas* V1442G in DCS^R^1, DCS^R^2, DCS^R^3, as well as the occurrence of *ppsA* E419K in the DCS^R^7, DCS^R^9, DCS^R^10 seem to have compensatory effects, alleviating any possible fitness cost associated with the *alr* D322N SNP (Fig. 2A). Furthermore, mutants DCS^R^8 and DCS^R^11 containing the promoter SNP did not exhibit any noticeable fitness cost over time. In line with previous mutants, DCS^R^8 and DCS^R^11 contain the additional SNP on *mas* V1442G that also compensates any associated fitness, as it did with the DCS^R^1, DCS^R^2, DCS^R^3 (Fig. 2A). The fact that there were no DCS resistant isolates obtained with only mutations in the promoter region of *alr* indicates that the original mutation might lead to a high fitness cost phenotype and thus either mutations in *mas* or *rv2962c* had to be established to provide compensatory effects and survival of these strains prior to promoter changes. DCS^R^7, DCS^R^9, DCS^R^10 seem to have a different compensatory mechanism with an additional mutation at the *ppsA* gene (Rv2931) E419K encoding for a polyketide synthase involved in the synthesis of phenolphthiocerol, also a part of pDIM molecules. The presence of two different compensatory mutations in two different genes involved in pDIM biosynthesis indicates that pDIM levels must have a protective effect, when *alr* activity is affected. Likely, these modifications are making *M. tuberculosis* further impermeable to DCS. Consistent with these findings, it has been found that mutations in the *ppsA* gene offer a fitness benefit in XDR strains (*36*). In addition, different mutations in the *ppsA*, *ppsB* and *ppsC* genes have been implicated in pyrazinoic acid resistance in *M. bovis* BCG (*37*). Although the mutations found by Gopal *et al*., and the ones described herein are not the same, none of our 11 DCS^R^ mutants showed any changes in the MIC_90_ to pyrazinamide or pyrazinoic acid (Fig. 1C and Fig. S3). Interestingly, our fitness kinetics experiments reveal that there was no significant fitness cost associated with mutations causing DCS resistance *in vitro*.

**Fig 2.**
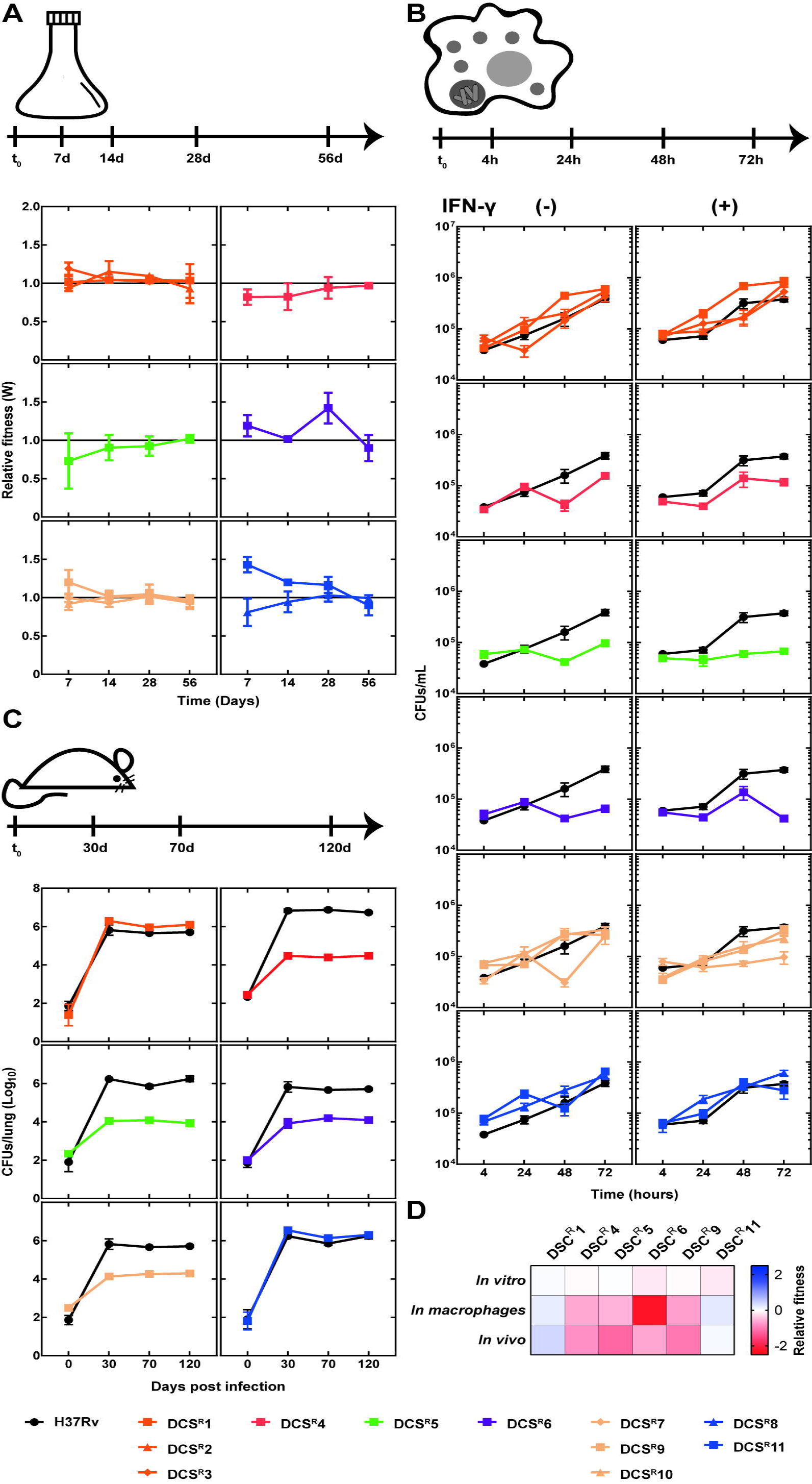
Fitness cost kinetics of DCS^R^ mutants. (**A**) Relative fitness of DCS^R^ mutants calculated using competition assays *in vitro*. DCS^R^ mutants are grouped according to their mutations and the relative fitness is followed over time. Parent strain relative fitness = 1. Data are representative of three independent experiments. (**B**) Relative fitness of DCS^R^ mutants obtained during macrophage infection. DCS^R^ mutants are grouped according to their mutations. Experiments were carried out with human blood-derived monocytes differentiated into macrophages. Bacterial uptake and survival inside naïve or interferon-gamma activated macrophages is compared over time with the wild type strain (black dots and line). Data shown is from one experiment representative of three independent experiments, where each macrophage culture originated from at least three blood donors and were pooled prior to infection to minimize donor variability. (**C**) Assessment of fitness cost associated with DCS resistance in the mouse model of infection. DCS^R^ mutants (1, 4, 5, 6, 9 and 11) and parent strain were used in a low-dose aerosol infection murine model (using C57BL/6 mice). The CFUs obtained from mouse lungs are shown for days 0, 30, 70 and 120 post-infection. Data for DCS^R^4 are representative of two independent experiments whereas for DCS^R^ mutants (1, 5, 6, 9 and 11) are from a single infection. Five mice were infected per strain per timepoint. (D). The relative fitness calculated for representative DCS^R^ mutants (1, 4, 5, 6, 9 and 11) at the endpoint of *in vitro* (day 56) competition assays, macrophages infection assays (day 3) and *in vivo* experiments (day 120), compared to the parent strain.

We then examined the uptake by and survival of the DCS^R^ strains in human monocyte-derived macrophages. All DCS^R^ strains tested were taken up similarly by macrophages (Fig. 2B). In line with the fitness cost shown *in vitro*, DCS^R^4, bearing solely the D322N Alr mutation, was attenuated during infection of naïve and of interferon gamma-activated human macrophages, compared to the parent strain (Fig. 2B). The presence of the *mas* V1442G or the *ppsA* E419K SNPs again led to compensatory benefits as strains DCS^R^1, DCS^R^2, DCS^R^3, DCS^R^7, DCS^R^9, DCS^R^8, DCS^R^10 and DCS^R^11 managed to infect, survive and proliferate inside macrophages at similar levels compared to the parent strain (Fig. 2B). In contrast, DCS^R^5, bearing the *alr* upstream deletion and a SNP on rv2962c gene was significantly attenuated for growth in human macrophages, compared to the parent strain (Fig. 2B). Similarly, DCS^R^6 also had an attenuated phenotype in human macrophages, indicating an associated fitness cost with Rv2962c T234I and Alr D322N mutations. These data suggest that some of the SNPs found in DCS^R^ strains are detrimental for the ability of *M. tuberculosis* to grow in human macrophages.

We next chose one DCS^R^ strain representative of each group for analysis of fitness/attenuation in the mouse model of infection. In line with the results obtained in Middlebrook medium and human macrophages, DCS^R^4 was significantly attenuated in the mouse model, compared to the parent strain, with a decrease in lung CFUs of more than two log_10_ during the chronic phase of infection (Fig. 2C). This failure to establish a high-burden infection in mice indicates that the Alr D322N mutation impairs *M. tuberculosis* cellular viability. The presence of the *mas* mutation (DCS^R^1) led to compensatory effects during the mouse infection as we observed before in medium and in macrophages. Unexpectedly, the presence of the *ppsA* mutation did not compensate *in vivo,* as seen with the mutant DCS^R^9 (Fig. 2C). No fitness compensation was observed with DCS^R^ strains having a mutation at the putative rhamnosyltransferase (Uniprot ID P9WN09), Rv2962c, including the *alr* repression mutant that was also found to be attenuated in mice (Fig. 2C). These results indicate that most of the DCS^R^ strains incur significant fitness cost *in vivo* that can be missed if only competition assays *in vitro* are used (Fig. 2D). In this case, clear compensatory mechanisms involving *mas* mutations can reverse the observed fitness costs associated with DCS resistance.

In order to understand the molecular and cellular mechanisms involved in DCS resistance in TB, we first investigated the gene expression pattern of a selected panel of genes including *alr* and *ddlA*, encoding both targets of DCS. The DCS^R^5, DCS^R^8 and DCS^R^11 mutants that contained either mutations in the *alr* promoter or a deletion upstream of *alr* overexpressed *alr* gene by more than 30-fold when compared to the parent strain (Fig.3A). This result indicates that the large deletion upstream *alr* found in DCS^R^5 likely removes a transcription factor binding site for an as yet unknown transcriptional repressor, but alternative mechanisms that lead to enhanced gene expression could also be involved. In contrast, all other DCS^R^ strains displayed similar levels of *alr* expression (Fig. 3A). There was no change in the expression of the *ddlA* gene in any strains indicating again that *ddlA* is not linked with DCS resistance in *M. tuberculosis*. Also, no changes in the RNA levels of genes related to downstream biochemical reactions in peptidoglycan biosynthesis such as *murD*, *murF* and *murI* were observed in any of the DCS^R^ strains (Fig. 3A). In addition, during an acute DCS treatment (high dose, short time), transcriptional changes of these selected genes were partially altered in the majority of the mutants. There is upregulation of DCS targets, *alr* and *ddlA,* as well as *cycA* and the downstream peptidoglycan biosynthesis related genes *murD*, *murF* and *murI* (Fig. 3A). These subtler transcriptional changes likely indicate a concerted effort to maintain the structural integrity of peptidoglycan in *M. tuberculosis* under DCS stress (Fig. 3A). We further examined the protein levels of Alr, using a specific anti-Alr serum, and found that Alr is also elevated in the DCS^R^5, DCS^R^8 and DCS^R^11 strains (Fig. 3B), consistent with the increase in *alr* RNA levels observed in Fig. 3A. Alr overexpression in DCS^R^5, DCS^R^8 and DCS^R^11 was also observed during acute exposure to DCS. Increased levels of Alr may serve the cell in two ways during DCS exposure: (i) providing extra D-Ala which will protect Ddl from DCS inhibition and (ii) removing free DCS from the cell, functioning as a “sponge”, a mechanism already shown in aminoglycoside resistance (*38*). Taken together, these results indicate that overexpression of Alr alone is sufficient to cause DCS resistance in *M. tuberculosis*.

**Fig. 3.**
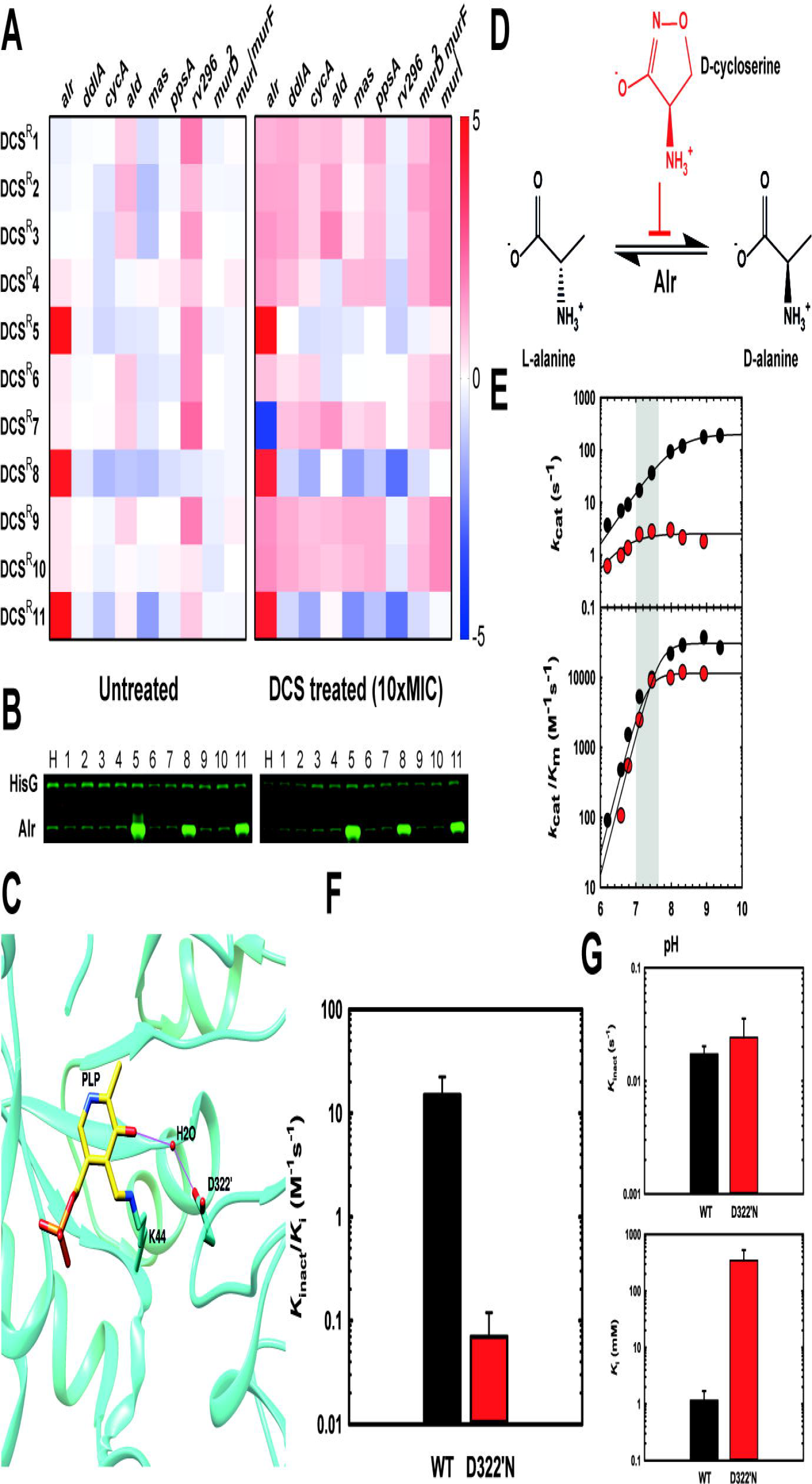
Cellular and molecular determinants of DCS resistance in *M. tuberculosis*. (**A**) qPCR results for differential gene expression (log_2_ transformed) of selected genes in the DCS^R^ mutants grown in the absence or presence of DCS (10 × MIC for 4h). Results are representative of three biological replicates. (**B**) Western-blot analysis of Alr expression in the DCS^R^ mutants and parent strain grown in the presence or absence of DCS (10 × MIC for 4h). HisG levels were used as loading control. Data are representative of two independent experiments. (**C**) Illustration of the active site of Alr from *M. tuberculosis* highlighting the location of residue D322 and its interactions with the pyridoxal 5’-phosphate cofactor through a water molecule-hydrogen bond network. (**D**) Reaction catalyzed by Alr and its inhibition by DCS. (**E**) pH-rate profile of the reaction catalyzed by wild-type and D322’N Alr. (**F**, **G** and **H**) Inactivation kinetics of wild-type and D322’N Alr by DCS.

We next investigated the effect of the D322N substitution on Alr activity, as this mutation was present in eight out of the eleven mutants isolated (~70% of the mutants). Analysis of the structure of *M. tuberculosis* Alr indicates that D322 from the adjacent monomer makes a water-mediated hydrogen bond with the pyridoxal 5’-phosphate cofactor (Fig. 3C). As such we will refer to it hereafter as D322’. Such an interaction suggests roles for this residue in either ligand binding, catalysis and/or inhibition. Recombinant Alr (wild-type) and Alr D322’N were expressed, purified and enzymatic activity as well as inhibition by DCS were evaluated (supplementary methods and Fig. 3D, E, F, G). We first analysed the pH profile of Ala racemization carried out by wild-type and the D322’N mutant. Under *k*_cat_ conditions, the D322’N mutant has significantly lower catalytic activity compared to wild-type Alr across all pH values tested (Fig.3E). At neutral pH, Alr D322’N is approximately 20-fold slower than wild-type enzyme. Under *k*_cat_/*K*_m_ conditions, no differences were observed for wild-type and D322’N Alrs in the pH range tested. As DCS forms a covalent inhibitor with alanine racemases pyridoxal 5’- phosphate cofactor and IC_50_ measurements are misleading for irreversible inhibitors, we evaluated *k*_inact_ and *K*_i_ parameters for wild-type and D322’N enzymes with DCS. The D322’N point mutation in Alr leads to a 240-fold decrease in the *k*_inact_/*K*_i_ for DCS (Fig. 3G). This result indicates that despite the moderate impact under *k*_cat_ conditions, Alr D322’N is highly resistant to DCS inhibition. Separation of *k*_inact_ (rate of inactivation) and *K*_i_ (binding) parameters revealed that the D322’N mutation leads to a loss of affinity for DCS (Fig. 3F), with only a very modest increase in the inactivation rate (Fig. 3F). Together, these results indicate that the most common mechanism to generate DCS resistance is by the modification of Alr, causing it to not bind effectively to DCS. Decreased affinity for DCS would lead to more alanine racemisation, which in turn would be predicted to protect Ddl from inhibition.

Prediction of resistance to novel antibiotics and pre-engineering/selection of compounds with ultra-low frequencies of resistance is essential for modern antibiotic development (*39*). Two more fundamental characteristics when evaluating drug resistance are the inherent rate to which mutations conferring resistance can arise in a particular pathogen and the potential ways in which fitness cost can be mitigated by alternative or compensatory mechanisms. The frequency of spontaneous mutations conferring resistance to DCS (mutation rate) is ultra-low in *M. tuberculosis* (ca. 10^-11^) ((*29*) and this work). Mutation rates are important but identifying and understanding potential routes for compensating fitness costs associated with resistance are equally important to evaluate and predict resistance appearance and dynamics in populations (*39*). Estimation of fitness costs not only in bacterial growth medium but also in cellular and whole-organism models is essential to uncover the true likelihood of a genotype to be propagated or eliminated from a population. We found that generation of mutants *in vitro* is an ideal way to access high variability of genotypes, which can then be studied in various models. This contrasts with the use of clinical strains, which have already been selected to allow growth in the host and therefore will only have low to negligible fitness costs. In addition, we found poor correlation between fitness measured in growth medium with fitness measured in macrophages and mice. *In vitro* estimations of bacterial fitness, which are more commonly carried out in a laboratory setting, seem to significantly underestimate phenotypes that are clear in cellular or whole-organism models.

This study is in agreement with studies highlighting the implication of *alr* in DCS resistance in *M. tuberculosis* (*22, 23*). We showed that there are two *alr*-related mechanisms underlying DCS resistance: overexpression of Alr or reduction of the inhibition of Alr through a previously unknown single point mutation. We showed that a promoter mutation upregulates *alr* gene transcription and increases Alr protein level up to 30-fold (Fig. 3A and B). This promoter mutation has been misannotated in recent clinical strains as a substitution of a Phe to Leu in position 4 of Alr; however, according to the recently curated version of Alr (Uniprot ID P9WQA9), this codon falls into the promoter sequence and it is not transcribed in the *alr* mRNA. In addition to this mutation, we also observed a deletion further upstream of the *alr* starting codon, possibly in an area where a yet-to-be-identified transcriptional repressor might bind. This deletion resulted in increased transcription of the *alr* gene with concomitant Alr overexpression, as observed in the promoter mutation strains. Crucially, this mutation has not been found in the clinic. The most common DCS resistance mechanism involves Alr modification (D322’N). This mutation reduces the affinity of Alr by DCS by over 240-fold, with some associated disruption of its catalytic activity. One XDR-TB strain contained a mutation on the succeeding residue, M221T, implicating the importance of these two residues in the inhibition of Alr by DCS.

In summary, DCS resistance is an ultra-rare event in *M. tuberculosis* and secondary compensatory mutations might alleviate potential fitness costs. Therefore, the lack of widespread clinical resistance to DCS is due to the low mutational frequency of *M. tuberculosis*. DCS is known to inhibit two enzymes from the same pathway and therefore it seems reasonable to propose that finding other molecules that truly inhibit more than one essential enzyme is the most rational route to resistance-evading antibiotics.

Lastly, the results of this work, in combination with clinical WGS studies and phenotypic data can help improve the diagnosis of DCS resistance through generation of molecular tools for mapping DCS-conferring mutations. Importantly, our findings can be utilized to improve the design of second-generation DCS analogues that will maintain low mutation rates and avoid resistance while minimizing undesirable cytotoxic effects.

## Acknowledgments

We thank Alex Gould, Kathy Niakan, Andreas Schaefer, Simon Waddell and Timothy McHugh for critical reading of the manuscript. We would like to thank the Crick’s Genomics/Equipment Park Science Technology Platform and the Media Preparation team for excellent technical assistance during this project. Thanks also to the Biological services staff at the Crick and NIBSC, particularly Sara Goulding, for assistance with animal welfare and maintenance. This work was performed under Home Office licences 70/8045 and P7611793C.

## Funding

This work was supported by the Francis Crick Institute which receives its core funding from Cancer Research UK (FC001060), the UK Medical Research Council (FC001060), and the Wellcome Trust (FC001060). TC was partially supported by the European Research Council (ERC) under the European Union’s Horizon 2020 research and innovation programme (grant agreement No 637730).

## Author contributions

GAP generated the DCS resistant mutants and carried out biochemical analysis of Alr and Alr-D322N. TC analyzed the whole genome sequencing data. DE performed Mtb experiments. AR, BMD and BK carried out mouse infection experiments. MMH contributed in implementation and oversight of mouse infection experiments at NIBSC. DE, GAP, MGG and LPSC designed, analyzed and interpreted data. DE wrote the first draft and all authors contributed during reviewing and editing of the final draft.

## Competing interests

Authors declare no competing interests.

## Data and materials availability

All data are available in the main text or the supplementary materials. Whole genome sequencing data of the strains used in this study is available in ArrayExpress (www.ebi.ac.uk/arrayexpress) under accession number E-MTAB-5935.

## Supplementary Materials

Materials and Methods

Figures S1-S3

Tables S1-S3

References (46-52)

